# Description of *Klebsiella africanensis* sp. nov., *Klebsiella variicola* subsp. *tropicalensis* subsp. nov. and *Klebsiella variicola* subsp. *variicola* subsp. nov

**DOI:** 10.1101/516278

**Authors:** Carla Rodrigues, Virginie Passet, Andriniaina Rakotondrasoa, Thierno Abdoulaye Diallo, Alexis Criscuolo, Sylvain Brisse

**Author notes:** These authors contributed equally to this work (Carla Rodrigues, Virginie Passet). “Correspondence and reprints”. **Abbreviations:** ANI, average nucleotide identity; isDDH, *in silico* DNA-DNA hybridization. E-mail addresses (C. Rodrigues), (V. Passet), (A. Rakotondrasoa), (T. A. Diallo), (A. Criscuolo).

## Abstract

The bacterial pathogen *Klebsiella pneumoniae* comprises several phylogenetic groups (Kp1 to Kp7), two of which (Kp5 and Kp7) have no taxonomic status. Here we show that group Kp5 is closely related to *Klebsiella variicola* (Kp3), with an average nucleotide identity (ANI) of 96.4%, and that group Kp7 has an ANI of 94.7% with Kp1 (*K. pneumoniae sensu stricto*). Biochemical characteristics and chromosomal beta-lactamase genes also distinguish groups Kp5 and Kp7 from other *Klebsiella* taxa. We propose the names *K. africanensis* for Kp7 (type strain, 200023^T^) and *K. variicola* subsp. *tropicalensis* for Kp5 (type strain, 1266^T^).

## 1. Introduction

The genus *Klebsiella* includes important human and animal pathogens and is largely distributed in animal carriage and the environment [1]. *Klebsiella pneumoniae* is the most common species isolated from infections of humans and other hosts. Brisse & Verhoef [2] reported the existence of *K. pneumoniae* phylogroups Kp1 to Kp4, and Blin *et al.* [3] reported two additional phylogroups, Kp5 and Kp6. Over the years, phylogenetic studies have demonstrated that *K. pneumoniae* phylogroups Kp1, Kp2, Kp4 and Kp3 correspond to different taxa, respectively *K. pneumoniae* (*sensu stricto*), *K. quasipneumoniae* subsp*. quasipneumoniae* and subsp*. similipneumoniae*, and *K. variicola* [4,5]. Phylogenetic analyses of Kp6 genomic sequences showed that this phylogroup corresponds to a recently proposed taxon, ‘*K. quasivariicola’* [6], which remains to be validly published. While screening for *K. pneumoniae* fecal carriage in Senegal, we identified strain 200023 (internal strain bank identifier, SB5857), which was not associated with any of the previous phylogroups or species, and which we here call Kp7. Together, *K. pneumoniae sensu stricto* and the other above referenced taxa and phylogroups constitute the *K. pneumoniae* complex. Typically, all members of this species complex are misidentified as *K. pneumoniae* or as *K. variicola* using standard laboratory methods, which masks their actual clinical and epidemiological significance [7–9]. The members of the *K. pneumoniae* complex can be correctly distinguished based on whole-genome sequencing (WGS) or by sequencing taxonomic marker genes such as *rpoB*. The chromosomal beta-lactamase gene also varies according to phylogroup: Kp1, Kp2, Kp3 and Kp4 harbor *bla*_SHV_, *bla*_OKP-A_, *bla*_LEN_ and *bla*_OKP-B_ beta-lactamase genes, respectively [10,11]. Recently, MALDI-TOF mass spectrometry protein biomarkers were described for the identification of *K. pneumoniae* complex members (Kp1 to Kp6) [12]. The aim of the present work was to define the taxonomic status of *K. pneumoniae* phylogroups Kp5 and Kp7.

## 2. Materials and methods

A total of thirty-seven strains of species, subspecies and phylogroups related to *K. pneumoniae* were included (Table 1, Fig. 1). Strain 1266^T^ (internal strain bank identifier, SB5531) was chosen as reference strain for phylogroup Kp5, whereas strain 200023^T^ (internal strain bank identifier, SB5857) was included as reference strain of Kp7 phylogroup. The genome (Accession Number ERS214332) of the strain 38679 [13] was included for phylogenetic comparisons even though the strain was not available at the time of this study. Strains were grown in tryptocasein soy agar (TSA) (BioRad, Marnes-La-Coquette, France). The biochemical characterization of the isolates was performed using API20E strips (BioMérieux) and Biolog phenotype microarrays plates PM1 and PM2 (Hayward, CA) following the protocol described by Blin *et al.* [3]. To extract the DNA, bacteria were plated on TSA and one colony was grown overnight with shaking at 37°C in 10 ml tryptocasein soy broth. DNA was extracted using the Wizard Genomic DNA purification kit (Promega, Charbonnières-les-Bains, France) and stored at −20°C. Genomic sequencing was performed using NextSeq-500 instrument (Illumina, San Diego, USA) with a 2 × 150 nt paired-end protocol.

**Table 1.**
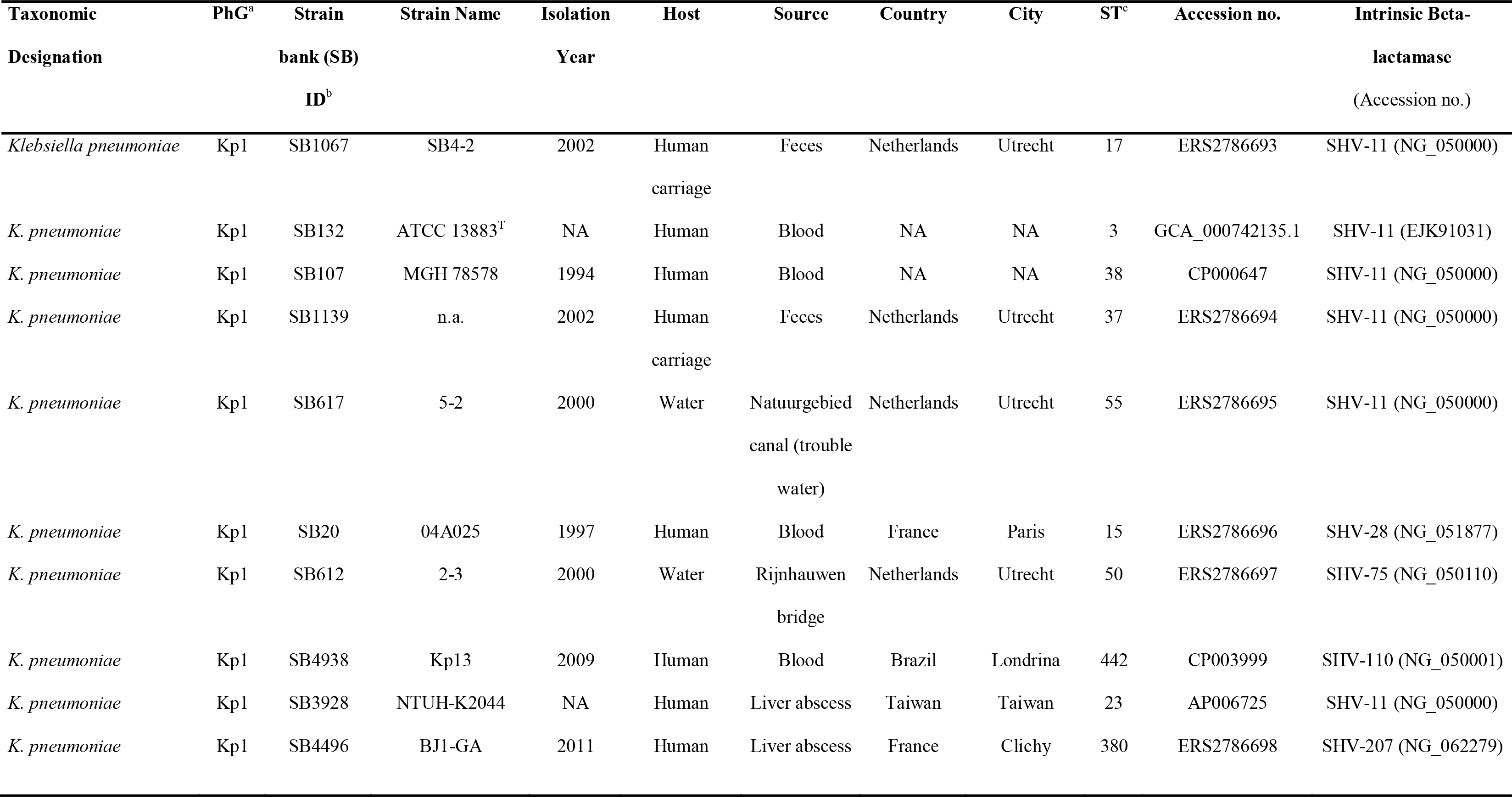
Strains included in this study

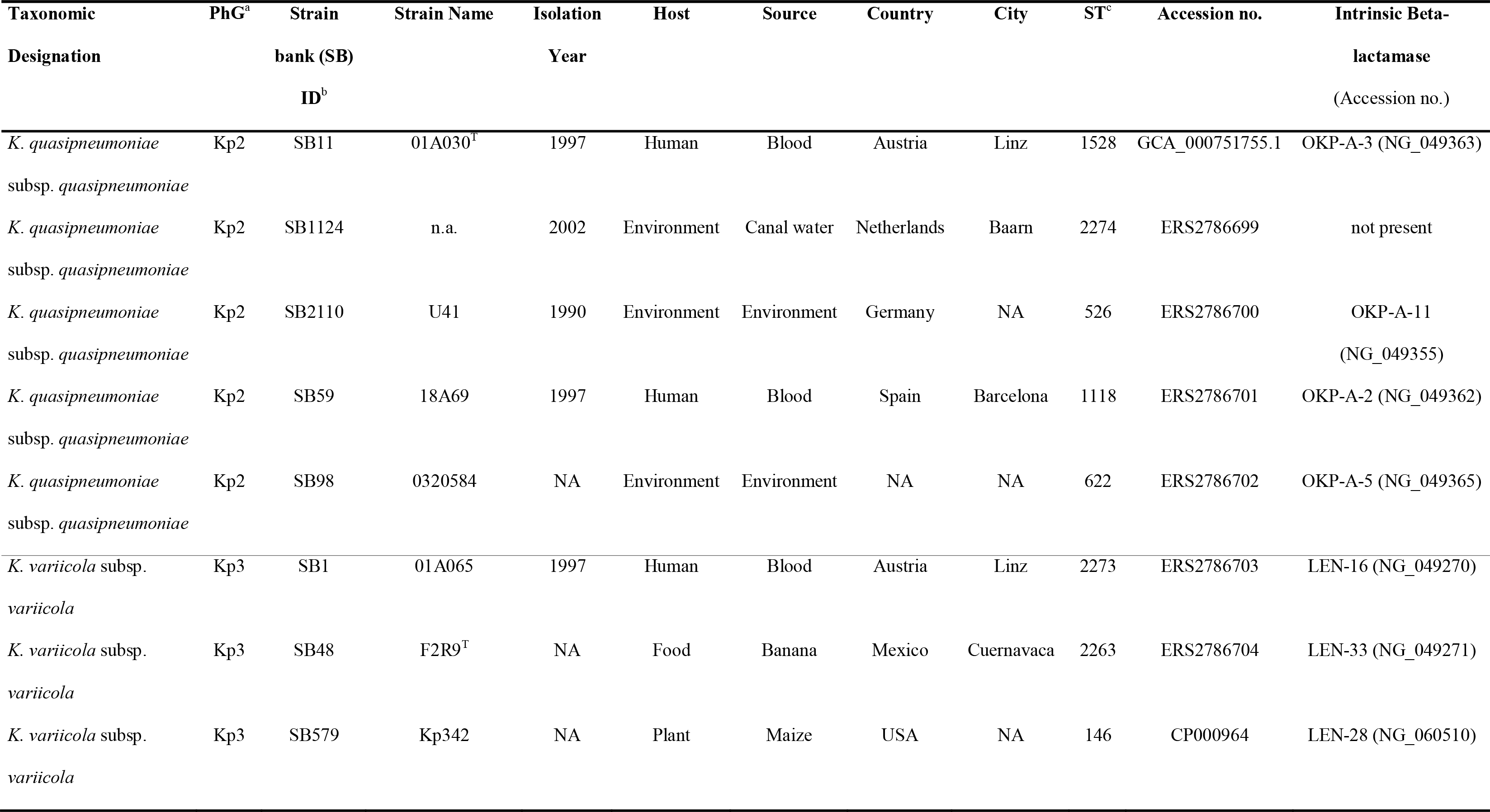

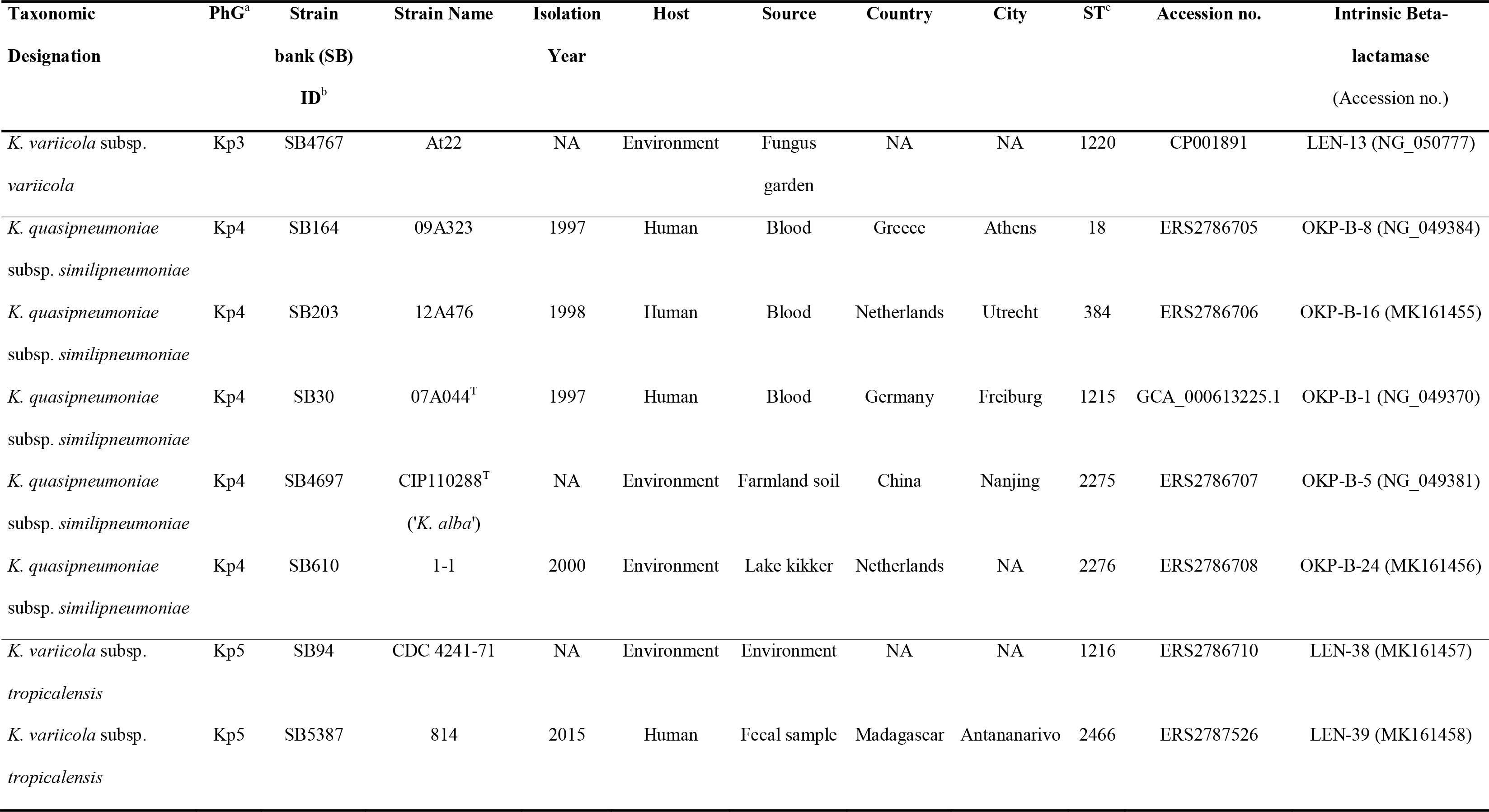

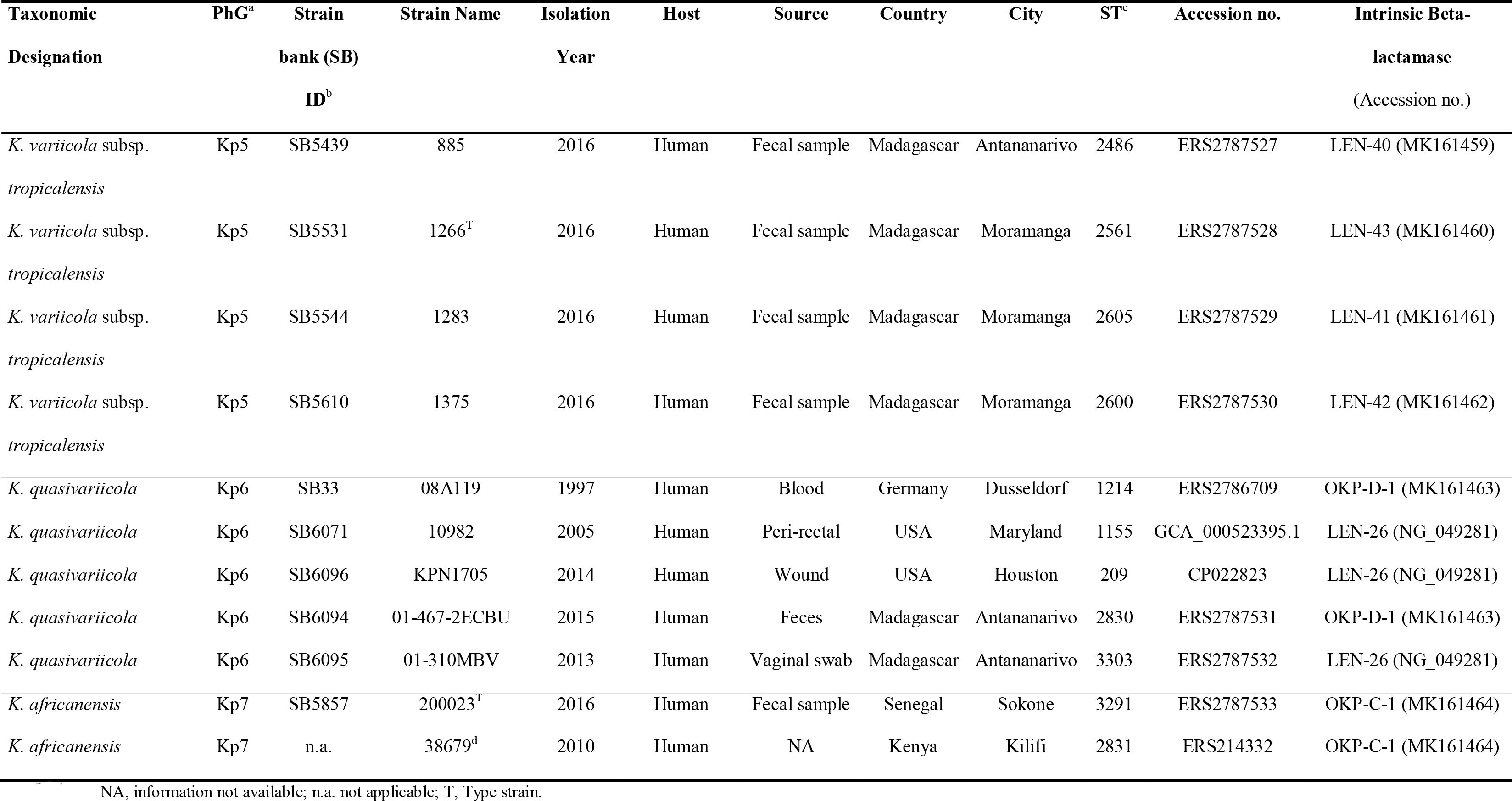

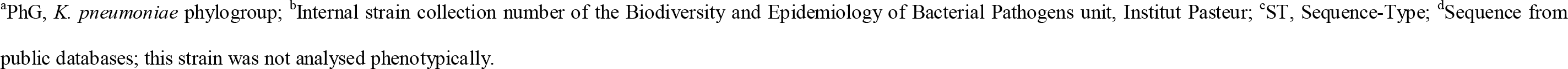

**Figure 1.**
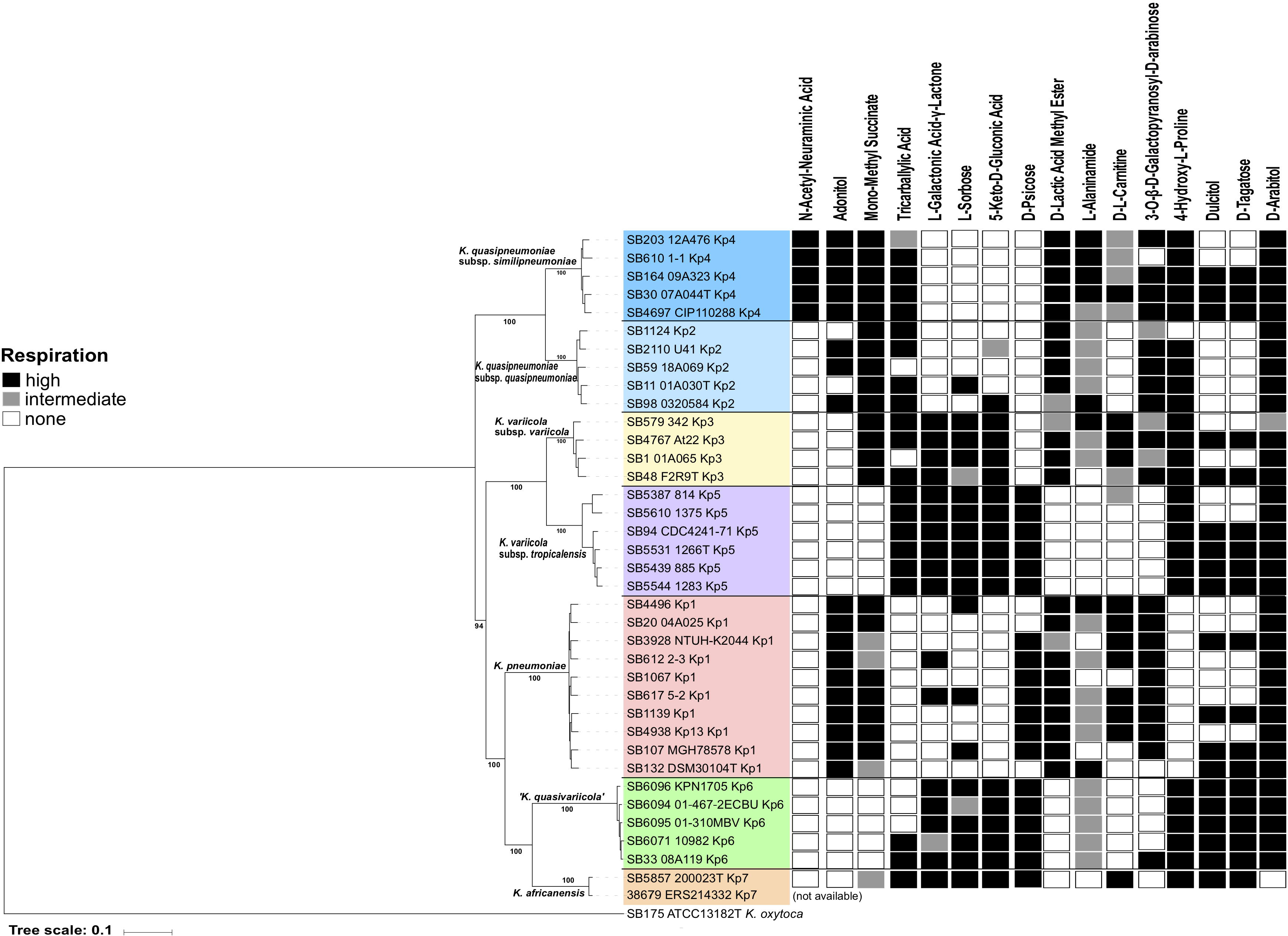
Maximum likelihood phylogeny and carbon sources utilization. **Left**: Phylogenetic tree (FastTree) based on the final recombination-free alignment of concatenated nucleotide sequence alignments of 1,703 core genes. Major taxonomic groups are indicated along the branches. Branch lengths represent the number of nucleotide substitutions per site (scale, 0.1 substitution per site). Bootstrap values are indicated at major nodes. **Right**: Metabolic phenotypes for the 16 most discriminant carbon sources. Black horizontal lines delimit phylogenetic lineages. Black squares correspond to substrate utilization; white square to absence of utilization; and grey squares to intermediate values. Strain 38679 (ERS214332) was not available for phenotyping.

Contig sequences were assembled using SPAdes v3.10.0 [14] and annotated with Prokka v1.12 [15]. JSpeciesWS [16] was used to calculate the BLAST-based average nucleotide identity (ANI). The *in silico* DNA-DNA hybridization (isDDH) was estimated using the GGDC tool (http://ggdc.dsmz.de; formula 2) [17]. Barrnap (https://github.com/tseemann/barrnap) was used to detect and extract 16S rRNA sequences from genome assemblies, whereas internal portions of genetic markers (*fusA*, *gapA*, *gyrA*, *leuS*, *rpoB*) were detected and extracted using BLAST and compared with previously reported sequences [4]. Chromosomal beta-lactamases were also extracted, and the new LEN and OKP amino-acid sequence variants were submitted to the Institut Pasteur MLST nomenclature database (https://bigsdb.pasteur.fr/klebsiella) for variant number attribution, and to NCBI for accession number attribution. Phylogenetic analyses based on the 16S rRNA gene sequence, concatenation of five housekeeping genes (2,609 aligned bp in total) and chromosomal beta-lactamases were performed using MEGA v7.0 with the neighbor-joining method. Genetic distances were estimated using the Jukes-Cantor substitution model in the case of nucleotide sequences and using the Jones-Taylor-Thornton (JTT) model for beta-lactamase protein sequences. For genome-based phylogenetic analysis, a phylogenetic tree was reconstructed from the concatenation of 1,703 genes defined as core genes using Roary v3.12 [18] with a BLASTP identity cut-off of 90%. *K. oxytoca* ATCC 13182^T^ (ERS2016112) was used as outgroup. Aligned characters affected by recombination events were detected and removed from the core gene alignment using Gubbins v2.2.0 [19]. A maximum-likelihood phylogenetic tree was inferred using FastTree v2.1.7 [20].

## 3. Results and Discussion

The nearly complete (1,462 nt) *rrs* gene sequence coding for 16S rRNA was compared among *K. pneumoniae* strains. Only a few sites showed reliable variation (Fig. S1), and all sequences were > 98.8% similar. The Kp5 reference strain 1266^T^ differed from F2R9^T^ (*K. variicola* type strain) by 6 (0.41%) nucleotide positions, whereas Kp7 strain 200023^T^ differed from KPN1705 (Kp6 reference strain) and DSM 30104^T^ (*K. pneumoniae* type strain) by 12 (0.82%) nucleotide positions in both cases. The phylogenetic information content of the highly conserved 16S rRNA sequence did not allow to reliably infer a phylogeny of 16S sequences (Fig. S1) as previously reported [21,22].

Maximum-likelihood phylogeny based on the recombination-purged nucleotide sequence alignments of 1,703 core genes (Fig. 1) showed seven highly supported branches. Phylogroup Kp5 was strongly associated with Kp3 (*K. variicola*), whereas Kp7 was associated with Kp6 and Kp1 (*K. pneumoniae*) groups. The latter result suggests that Kp6 and Kp7 share a common ancestor with *K. pneumoniae* and may therefore represent the closest relatives of the clinically most significant member of the species complex. Phylogenies based on the five concatenated housekeeping genes *fusA*, *gapA*, *gyrA*, *leuS* and *rpoB*, and of chromosomal beta-lactamases, both corroborated the distinction among the seven groups (Fig. S2 and Fig. S3). The phylogeny of chromosomal beta-lactamases was highly concordant with core genes phylogeny, with the remarkable exception of phylogroup Kp6, which harbored two different types of beta-lactamases, *bla*_LEN_ and *bla*_OKP-D_, depending on the strain. This is consistent with frequent homologous recombination observed in the Kp6 phylogroup using Gubbins (data not shown) and as previously reported [23]. In order to estimate the genome-wide divergence among phylogroups, average nucleotide identity (ANI) was used. The ANI values of Kp5 (strain 1266^T^) with *K. variicola* F2R9^T^ (Kp3) was 96.4%, above the cut-off value (approximately 95-96%) proposed for species distinction [24,25]. In contrast, the ANI values of Kp7 strain 200023^T^ with ‘*K. quasivariicola’* KPN1705 (Kp6) and *K. pneumoniae* DSM30104^T^ (Kp1) were 94.9% and 94.7%, respectively (Table 2). The isDDH relatedness of 1266^T^ (Kp5) with Kp3 strains ranged from 75.0% to 75.5%, but ranged from 51.2% and 56.0% with the other phylogroups. For 200023^T^ the isDDH relatedness with Kp1 to Kp6 phylogroups was between 50.5% and 61.9%, well below the ~70% cut-off used to define bacterial species. Therefore, both ANI and isDDH analyses lead us to consider that 1266^T^ is a subspecies of *K. variicola* (Kp3) and that 200023^T^ represents a novel *Klebsiella* species.

**Table 2.** Average nucleotide identity (ANI) values obtained among seven members of the *Klebsiella pneumoniae* complex

| Query genome<sup>a</sup> | Size (nucleotides) | DNA G+C content<br>(mol%) | Average nucleotide identity of test genome against query genomes |  |  |  |  |  |  |
| --- | --- | --- | --- | --- | --- | --- | --- | --- | --- |
|  |  |  | Kp1 | Kp2 | Kp3 | Kp4 | Kp5 | Kp6 | Kp7 |
| **Kp1** | 5 234 536 | 57.3 | * | 93.21 | 94.14 | 93.40 | 93.47 | 93.51 | 94.69 |
| **Kp2** | 5 343 479 | 58.1 | 93.21 | * | 93.06 | 96.27 | 92.80 | 92.36 | 92.64 |
| **Kp3** | 5 470 009 | 57.6 | 94.03 | 92.94 | * | 93.23 | 96.65 | 93.49 | 93.66 |
| **Kp4** | 5 084 979 | 58.3 | 93.49 | 96.33 | 93.42 | * | 93.03 | 92.52 | 92.82 |
| **Kp5** | 5 698 764 | 57.1 | 93.22 | 92.40 | 96.38 | 92.69 | * | 93.00 | 93.03 |
| **Kp6** | 5 540 188 | 56.9 | 93.50 | 92.37 | 93.58 | 92.50 | 93.27 | * | 94.85 |
| **Kp7** | 5 156 720 | 57.3 | 94.66 | 92.58 | 93.75 | 92.66 | 93.31 | 94.86 | * |

<sup>a</sup>Kp1, *K. pneumoniae* DSM30104<sup>T</sup>; Kp2, *K. quasipneumoniae* subsp. *quasipneumoniae* 01A030<sup>T</sup>; Kp3, *K. variicola* subsp. *variicola* F2R9<sup>T</sup>; Kp4, *K. quasipneumoniae* subsp. *similipneumoniae* 07A044<sup>T</sup>; Kp5, *K. variicola* subsp. *tropicalensis* 1266<sup>T</sup>; Kp6, *K. quasivariicola* KPN1705; Kp7, *K. africanensis* 200023<sup>T</sup>

The phenotypic characteristics of the strains were compared (Table S1, Fig. 1). Using API20E strips we confirmed that all Kp5 strains and 200023^T^ were positive for the utilization of lactose and mannitol, for the urease, malonate, lysine decarboxylase, Voges–Proskauer and ONPG tests, and reduced nitrate to nitrite, whereas they were negative for indole and ornithine decarboxylase. Furthermore, microscopy confirmed that all strains were non-motile. The use of phenotype microarrays provided insights into the metabolic profiles of the seven phylogroups and identified carbon sources that differentiate Kp5 and Kp7 strains form other groups (Fig. 1). The inability to metabolize mono-methyl succinate, D-lactic acid methyl ester and 3-O-(ß-D-galactopyranosyl)-D-arabinose, and the ability to metabolize D-psicose differentiated Kp5 strains from *K. variicola*. Kp7 strain 200023^T^ was unique in being unable to metabolize D-arabitol and was also differentiated from ‘*K. quasivariicola*’ and *K. pneumoniae*, its closest phylogenetic neighbors, by the ability to metabolize D-L-carnitine and 4-hydroxy-L-proline, respectively.

Based on the above genomic and phenotypic characteristics, we propose that Kp5 be considered as a subspecies of *K. variicola*, which we propose to name *K. variicola* subsp. *tropicalensis*; whereas Kp7 represents a novel species, which we propose to name *K. africanensis*.

### 3.1 Description of *Klebsiella variicola* subsp. *tropicalensis* subsp. nov

(tro.pi.cal.en’sis. N.L. fem. adj. *tropicalensis*, referring to the tropical area of Earth, from where Kp5 strains have been isolated).

The description is based on six strains. Cells are gram negative, non-motile, non-spore forming, straight, rod-shaped, capsulated. Colonies are smooth, circular, white, dome-shaped, glistening. The general characteristics are as described for *K. pneumoniae*. Urease positive, ONPG positive, Voges-Proskauer test positive, indole negative. Lysine decarboxylase positive, ornithine decarboxylase negative. Distinguished from the other *K. pneumoniae* complex members by the characteristics listed in Table S1. *K. variicola* subsp. *tropicalensis* strains are able to metabolize tricarballylic acid, L-galactonic acid-γ-lactone, L-sorbose, 5-keto-D-gluconic acid, D-psicose, 4-hydroxyl-L-proline and D-arabitol. They are not able to metabolize N-acetyl-neuraminic acid, adonitol, mono-methyl succinate, D-lactic acid methyl ester, L-alaninamide and 3-O-b-galactopyranosyl-D-arabinose. *K. variicola* subsp. *tropicalensis* isolates were recovered from asymptomatic fecal carriage and from the environment.

The type strain is strain 1266^T^ (= SB5531^T^, {CIP and DSMZ numbers: *in process*}), isolated in 2016 from a fecal sample of a human asymptomatic carrier in Antananarivo, Madagascar. The EMBL (GenBank/DDBJ) accession numbers of the *rrs* (coding for 16S rRNA) gene is MK040621. The genome sequence accession number is ERS2787528.

### 3.2 Description of *Klebsiella variicola* subsp.*variicola* subsp. nov

(va.ri.i’co.la. adj. varius different, differing, various, L. suffix n. -cola inhabitant, N.L. fem./masc. n. *variicola* inhabitant of different places)

In accordance to Rule 46 of the Bacteriological Code, the description of *Klebsiella variicola* subsp *. tropicalensis* automatically creates a second subspecies, *Klebsiella variicola* subsp *. variicola* subsp. nov. for which the type strain is F2R9^T^ (= ATCC BAA-830^T^, DSM 15968^T^) [6]. In addition to the phenotypic characters consistent within all members of the *K. pneumoniae* complex, reactions common to both subspecies of *K*. *variicola* are the ability to metabolize tricarballylic acid, L-galactonic acid-γ-lactone, L-sorbose, 5-keto-D-gluconic acid, 4-hydroxyl-L-proline and D-arabitol, and the inability to ferment adonitol and N-acetyl-neuraminic acid. The ability to metabolize mono-methyl succinate, D-lactic acid methyl ester and 3-O-(ß-D-galactopyranosyl)-D-arabinose, and the inability to metabolize D-psicose differentiated *K. variicola* subsp. *variicola* from *K. variicola subsp. tropicalensis*.

### 3.3 Description of *Klebsiella africanensis* sp. nov

(a.fri.can.en’sis. N.L. fem. adj. *africanensis*, referring to Africa, from where Kp7 strains have been isolated).

The phenotypic description is based on one strain. Cells are gram negative, non-motile, non-spore forming, straight, rod-shaped, capsulated. Colonies are smooth, circular, white, dome-shaped, glistening. The general characteristics are as described for *K. pneumoniae*. Urease positive, ONPG positive, Voges-Proskauer test positive, indole negative. Lysine decarboxylase positive, ornithine decarboxylase negative. Distinguished from the other *K. pneumoniae* complex members by the characteristics listed in Table S1. Distinguished from all the other *K. pneumoniae* complex members by the inability to metabolize D-arabitol. *K. africanensis* type strain is able to metabolize tricarballylic acid, L-galactonic acid-γ-lactone, L-sorbose, 5-keto-D-gluconic acid, D-psicose, L-carnitine, 4-hydroxyl-L-proline, D-dulcitol, and D-tagatose. Is not able to metabolize N-acethyl-neuraminic acid, adonitol, D-lactic acid methyl ester, L-alaninamide and 3-O-b-galactopyranosyl-D-arabinose and D-arabitol. So far, *K. africanensis* was recovered from asymptomatic human fecal carriage in Senegal (this study) and from human clinical samples in Kenya [13].

The type strain is strain 200023^T^ (= SB5857^T^, {CIP and DSMZ numbers: *in process*}), isolated in 2016 from a fecal sample of a human asymptomatic carrier in Sokone, Senegal. The EMBL (GenBank/DDBJ) accession numbers of the *rrs* (coding for 16S rRNA) gene is MK040622. The genome sequence accession number is ERS2787533.

### 3.4 Nucleotide sequence accession numbers

The genomic sequences of strains 1266^T^ and 200023^T^ were submitted to the European Nucleotide Archive under accession numbers ERS2787528 and ERS2787533, respectively. 16S rRNA sequences were also submitted individually under the accession numbers MK040621 and MK040622, respectively. The genome sequence data of strains 814, 885, 1283, 1375, 01-467-2ECBU and 01-310MBV were submitted to the European Nucleotide Archive under accession numbers ERS2787526, ERS2787527 and ERS2787529 to ERS2787532, respectively (under the BioProject number PRJEB29143).

## Acknowledgements

We thank the ‘P2M-Plateforme de Microbiologie Mutualisée’ (PIBnet) from Institut Pasteur for genomic sequencing. We acknowledge Jean-Marc Collard (Institut Pasteur of Madagascar), Bich-Tram Huynh (Institut Pasteur, France) and Raymond Bercion (Institut Pasteur of Dakar) for support in *K. pneumoniae* carriage studies. We also thank to S. Wesley Long (Houston Methodist Hospital) and David A. Rasko and J. Kristie Johnson (University of Maryland School of Medicine, Baltimore) for providing strains analyzed in this study.

---

**Figure S1.** Phylogenetic relationships (neighbor-joining method, Jukes-Cantor correction) based on the sequence of the *rrs* gene coding for 16S rRNA and the respective multiple sequence alignment restricted to the variable positions.

The tree was rooted using *K. oxytoca* ATCC 13182^T^ sequence. Bootstrap proportions obtained after 1000 replicates are indicated at the nodes. Branch lengths represent the number of nucleotide substitutions per site (scale, 0.001 substitution per site). Strain labels are given as Strain Bank ID (*e.g*., SB11) followed by original strain name, followed by phylogroup. A ‘T’ after the strain name indicates type strains.

**Figure S2.** Phylogenetic relationships (neighbor-joining method, Jukes-Cantor correction) based on the concatenated sequences of the five-individual protein-coding genes *fusA*, *gapA gyrA, leuS* and *rpoB*.

The was rooted using *K. oxytoca* ATCC 13182^T^. Bootstrap proportions obtained after 1000 replicates are indicated at the nodes. Branch lengths represent the number of nucleotide substitutions per site (scale, 0.01 substitution per site). Strain labels are given as Strain Bank ID (*e.g*., SB11) followed by original strain name, followed by phylogroup. A ‘T’ after the strain name indicates type strains.

**Figure S3.** Phylogenetic tree based on the core gene nucleotide sequences (A, maximum likelihood) compared to the gene tree of the chromosomal beta-lactamase amino-acid sequences (B, neighbor-joining method, Jones-Taylor-Thornton model).

Branch lengths represent the number of nucleotide (A) or amino-acid (B) substitutions per site (scale, 0.1 substitution per site). Major taxonomic groups are indicated along the branches of the phylogenetic tree A (*Kqs*, *K. quasipneumoniae subsp*. *similipneumoniae; Kqq*, *K. quasipneumoniae* subsp. *quasipneumoniae*, *Kvv*, *K. variicola* subsp. *variicola*, *Kvt*, *K. variicola* subsp. *tropicalensis; Kpn*, *K. pneumoniae*; *Kqv*, ‘*K. quasivariicola’*, *Kaf*, *K. africanensis*). Strain labels are given as Strain Bank ID (*e.g*., SB11) followed by original strain name, followed by phylogroup. A ‘T’ after the strain name indicates type strains. In panel B the chromosomal beta-lactamase and the corresponding accession number are indicated beside the strain names. Node labels indicate bootstrap values based on 1,000 replicates. Red stars indicate remarkable discrepancies between the two phylogenetic trees (phylogroup Kp6 presents two types of chromosomal beta-lactamases).

