## Supplementary figures and images for "Description of *Klebsiella africanensis* sp. nov., *Klebsiella variicola* subsp. *tropicalensis* subsp. nov. and *Klebsiella variicola* subsp. *variicola* subsp. nov"

### Figure S1

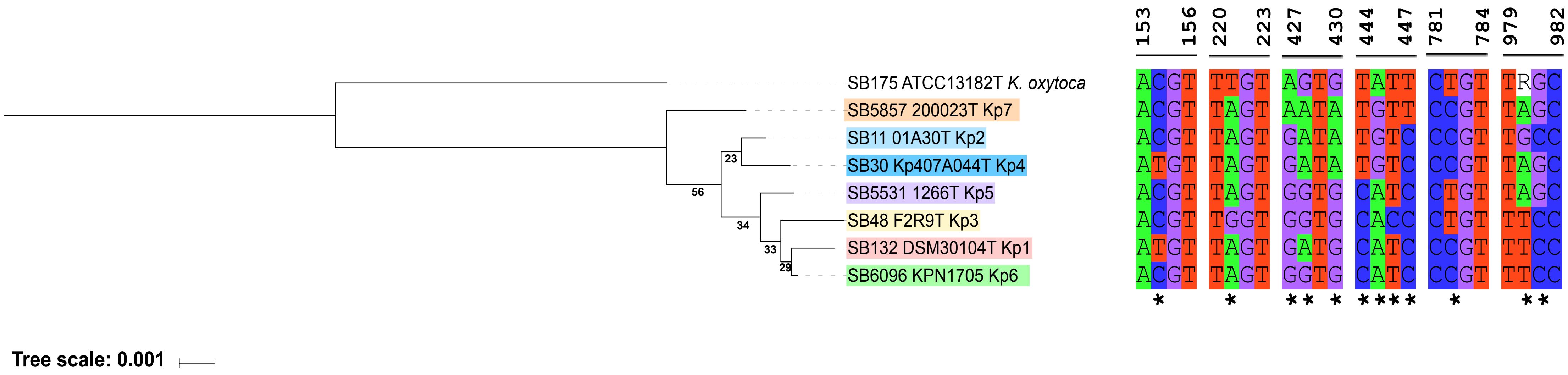

### Figure S2

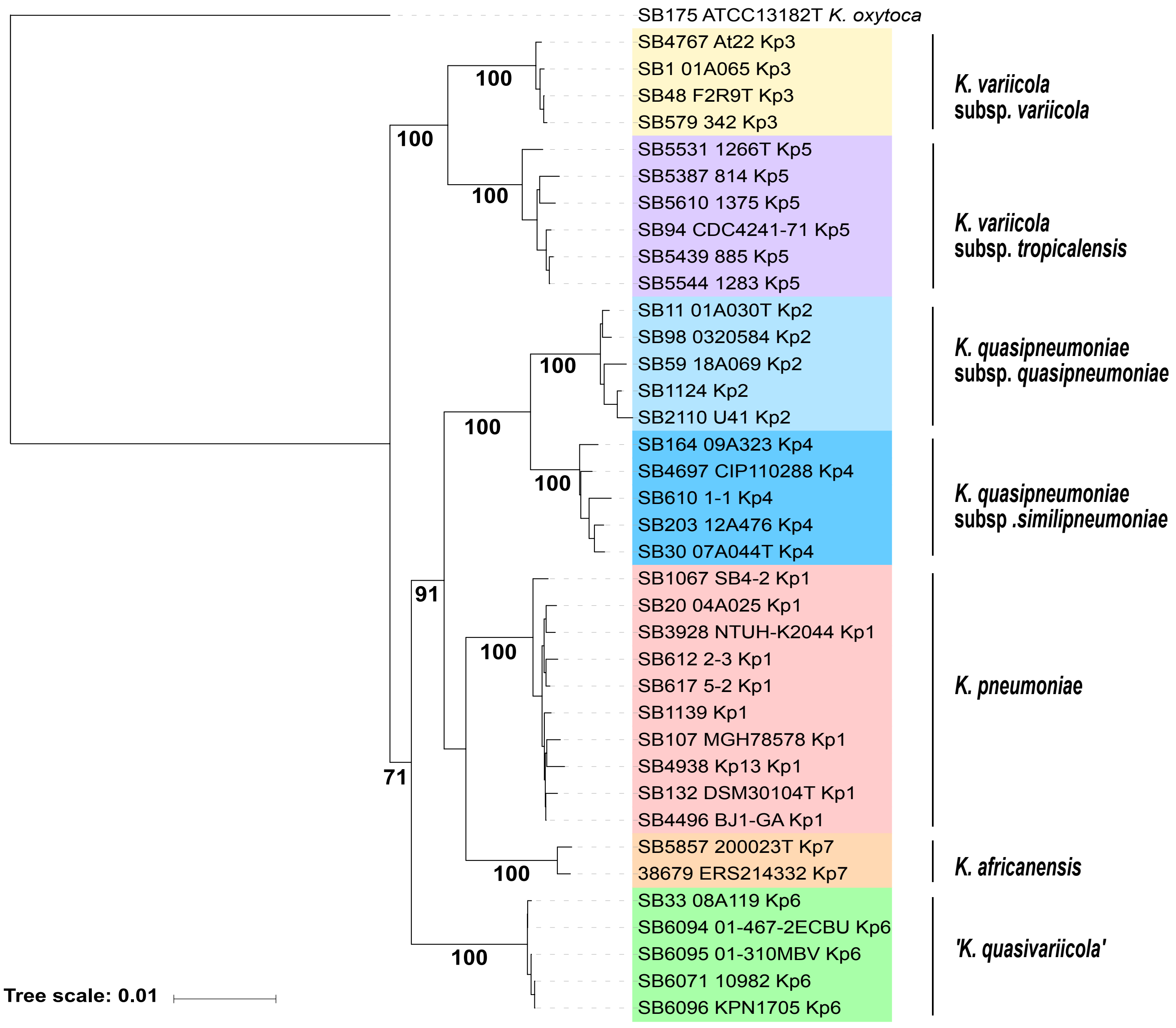

### Figure S3

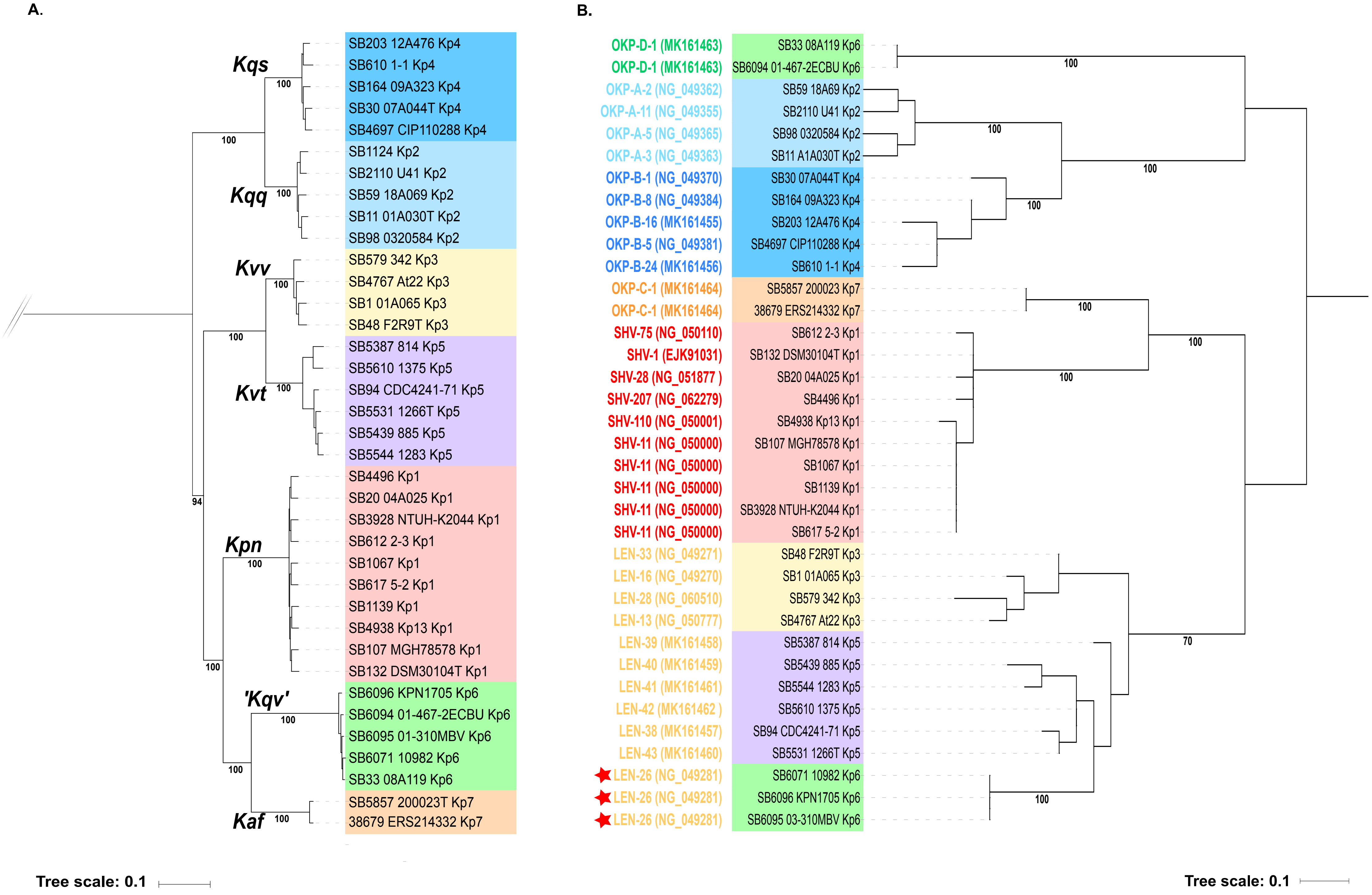
