## Supplementary material for "Description of *Klebsiella africanensis* sp. nov., *Klebsiella variicola* subsp. *tropicalensis* subsp. nov. and *Klebsiella variicola* subsp. *variicola* subsp. nov": Table S1

**Table S1**. Differential characteristics of the taxa under study

|  | ***K. pneumoniae* phylogroup** ^a^ (No. of strains) | | | | | | |
| --- | --- | --- | --- | --- | --- | --- | --- |
|  | **Kp1** (n=10) | **Kp2** (n=5) | **Kp4** (n=5) | **Kp3** (n=4) | **Kp5** (n=6) | **Kp6** (n=5) | **Kp7** (n=1) |
| **Metabolic phenotypes** |  |  |  |  |  |  |  |
| Dulcitol | v | - | v | v | v | + | + |
| Adonitol | + | v | + | - | - | - | - |
| Tricarballylic acid | - | + | + | v | + | v | + |
| Mono methyl succinate | v | + | + | + | - | - | + |
| D-psicose | v | - | - | - | + | + | + |
| L-galactonic acid-γ-lactone | - | - | - | + | + | + | + |
| N-acetyl-neuraminic acid | - | - | + | - | - | - | - |
| D-arabitol | + | + | + | + | + | + | - |
| 3-O-(ß-D-galactopyranosyl)-D-arabinose | + | + | + | + | - | - | - |
| L-sorbose | v | - | - | + | + | + | + |
| D-tagatose | v | - | v | v | v | + | + |
| 5-keto-D-gluconic acid | - | v | - | + | + | + | + |
| D-lactic acid methyl ester | + | + | + | + | - | - | - |
| L-alaninamide | v | + | + | v | - | + | - |
| 4-hydroxyl-L-proline | - | + | + | + | + | + | + |
| L-carnitine | v | - | + | + | + | - | + |

-, less than 20% of strains positive; +, more than 80% of strains positive; v, between 20% and 80% of strains positive.

^a^Kp1, *K. pneumoniae*; Kp2, *K*. *quasipneumoniae* subsp. *quasipneumoniae*; Kp3, *K. variicola* subsp. *variicola*; Kp4, *K. quasipneumoniae* subsp. *similipneumoniae*; Kp5, *K. variicola* subsp. *tropicalensis*; Kp6, *K. quasivariicola*; Kp7, *K. africanensis.*
